# Oxytocin promotes lying for personal gain in a genotype-dependent manner

**DOI:** 10.1101/361212

**Authors:** Cornelia Sindermann, Ruixue Luo, Benjamin Becker, Keith M Kendrick, Christian Montag

## Abstract

Society values honesty, since it fosters trust in others. Although we have a strong moral aversion to lying, particularly when it is self-serving, we nevertheless lie quite frequently and the biological basis for this is poorly understood. The hypothalamic neuropeptide oxytocin has been implicated in a number of anti-social as well as pro-social behaviours, including lying to benefit in-group members or in competitive situations. The aim of the present study was to investigate the effects of oxytocin administration on self-serving lying behaviour and possible moderating effects of genetic underpinnings of the oxytocin receptor. A total of 161 adult men participated in a randomized double-blind placebo-controlled between-subject intranasal oxytocin administration (24 International Units) study where self-serving lying was assessed using the die-in-a-cup paradigm. Additionally, contributions of polymorphisms in the oxytocin receptor gene were investigated using a haplotype approach. Results showed that while placebo-treated subjects behaved honestly across three successive rounds, oxytocin administration promoted self-serving lying, particularly in the third / last round and only to a certain degree (not to the maximum). Moreover, this effect of oxytocin was strongest in carriers of the GCG individual haplotype (rs237887-rs2268491-rs2254298) and non-carriers of the GT individual haplotype (rs53576-rs2268498) on the oxytocin receptor gene. Overall our findings demonstrate that oxytocin administration can promote self-serving lying when subjects are given repeated opportunities to lie and that these effects are moderated by genetic underpinnings of the oxytocin receptor.

## Introduction

While honesty is an important moral behaviour (see for example [1, 2]) and many people claim for themselves to be honest (e.g. [3]), lying is still prevalent and often generates public indignation and criticism, such as Bill Clinton denying an affair with Monica Lewinsky and Bernie Madoff causing billions of dollars of costs due to a ponzi scheme fraud. Results of a diary study indicate that also “normal” people lie in around 20-33% of their everyday social interactions [4]. In this context it is important to note that different kinds of lies exist and that reasons to lie are manifold [4]; hence, not all lies are considered as immoral or socially un-acceptable [5]. But lying to enhance one`s own payoffs or reduce one’s own costs, namely self-serving lying, is particularly seen as immoral and a violation of social norms since it can disrupt social relations by damaging interpersonal trust or results in a cost for others.

Despite the frequent occurrence of (self-serving) lying in everyday life, surprisingly little is known about its biological underpinnings. Two previous studies have reported that intranasal administration of the hypothalamic neuropeptide oxytocin (OXT) increases lying for financial gain when it benefits an in-group (including oneself) [6], or in a competitive situation in association with conformity to perceived deceptiveness of others [7]. However, using a coin-toss (prediction) task in which participants have only two possible choices – to “lie” or “be honest” –, neither study found evidence for effects on pure self-serving lying. Dishonesty is however subject to different gradations and this can be investigated using the so-called “die-in-a-cup” paradigm where subjects have the opportunity to lie to varying degrees [8, 9]. This paradigm has, for example, been used to demonstrate that testosterone administration reduced lying in males [10] and testosterone often shows opposing effects compared to OXT [11]. Additionally, previous studies have not addressed the question of whether OXT effects might vary with increased number of opportunities to lie. Clearly, the decision to exhibit dishonesty can change when there is a repeated chance to lie, dependent upon the actual outcome in previous rounds (lucky or unlucky) and the extent to which participants are convinced that lying would be detected and/or punished. Thus, OXT might, for example, only promote self-serving lying when subjects are completely convinced that their lies will not be detected and the level of certainty is likely to increase with every successful round of lying.

In addition, variations in the oxytocin receptor (OXTR) gene may influence the propensity for self-serving lying. Heritability estimates of lying are around 29-42% [12]. Also, the impact of intranasal OXT administration on self-serving lying might be moderated by genetic underpinnings of the OXTR. Support comes from studies reporting moderating effects of polymorphisms in the OXTR gene, located at chromosome 3p25.3 [13], on OXT administration effects on various measures of social behaviour and cognition [14-16]. But to date no study has investigated such effects on self-serving lying.

To this end the present psycho-pharmaco-genetic study aimed at investigating the interaction effects between variations in the OXTR gene and the intranasal administration of OXT on self-serving lying when participants are given the chance to lie repeatedly without negative consequences using repeated rounds of the die-in-a-cup paradigm. Given the non-significant findings on self-serving lying in previous OXT studies, but also taking into account the differences between the tasks of the studies, it was hypothesized that i) the tendency towards lying will be influenced by intranasal OXT administration ii) the effect might be in particular visible when participants have repeated opportunities to lie undetectably, iii) this effect will be moderated by genetic variation(s) of the OXTR gene.

## Materials and Methods

### Participants

Participants were first recruited for the Chengdu Gene Brain Behaviour Project (CGBBP) where they provided buccal cells for genotyping as well as completed a number of questionnaires including the HEXACO-PI-R Honesty-Humility scale [17]. After participation in the CGBBP, male participants were invited to participate in a randomized double-blind placebo-controlled between-subject intranasal OXT administration study. Exclusion criteria were any contraindication for OXT administration (e. g. hypersensitivity to OXT, nasal congestion), neurological or psychiatric disorders (including drug/alcohol abuse), regular or current medication and participation in another OXT administration study within the last 6 months prior to the present experiment. In total N=176 Chinese males (M_age_=21.33, SD=2.48) participated in the present experimental study. Participants were asked to sleep as usual on the day before testing and to abstain from caffeine-containing beverages on the day of the experiment. Due to technical failures, missing data and/or as a result of misunderstood instructions, n = 15 participants (8 receiving PLC, 7 receiving OXT) were excluded from the final analysis leading to a final sample-size of N=161 participants (n=80 PLC, n=81 OXT; M_age_=21.12, SD=2.50; n=149 Han). The study was approved by the local ethics committee at the University of Electronic Science and Technology of China (UESTC) Chengdu, China. Procedures were in accordance with the latest revision of the Declaration of Helsinki. All participants gave written informed consents prior to participation in both the CGBBP and the present experimental study.

### Genotyping and haplotype analyses

Detailed information about genotyping and haplotype analyses as well as distributions and (sub-)sample sizes of genotypes and individual haplotypes are provided in the Supplementary Material. Five OXTR SNPs were investigated in each participant and within these two haplotype blocks could be identified comprising i) OXTR rs237887-rs2268491-rs2254298, ii) rs53576-rs2268498. In each haplotype block, three individual haplotypes were investigated: GCG-, GTA-, ACG- (rs237887-rs2268491-rs2254298), AC-, AT- and GT- (rs53576-rs2268498). For each of the individual haplotype groups of carriers vs. non-carriers were built.

### Experimental procedure

#### Oxytocin-challenge study

For participation in the randomized double-blind placebo-controlled between-subject design experimental study each participant received a basic payment plus the money they individually earned in the economic games (participants also took part in a dictator and an ultimatum game after the die-in-a-cup paradigm. Only the die-in-a-cup paradigm results are presented here since the other two paradigms have different objectives and involve interactions with others). Each participant sat in a separated (obscured) cubicle. Hence, neither the experimenters nor the participants could see each other during the experiments. After arriving in the laboratory participants first filled in some basic demographic information. Afterwards, the self-administration of OXT (24 International Units) was implemented under the supervision of the trained experimenters. The complete procedure was in accordance with standardized guidelines [18] and detailed information is provided in the Supplementary Material. The participants were not able to guess better than chance if they retrieved PLC or OXT (Chi^2^(1)=2.30 p=.129 (N=161)), confirming successful double-blinding. The experiment was carried out blinded for the genetic data.

#### The die-in-a-cup paradigm

The die-in-a-cup paradigm (similar to the procedure used in for example [8-10]) was explained by standardized on-screen presentations (in case of problems or questions, the experimenters were available). The participants were first informed that they would receive an additional payoff for the following task according to the numbers they threw on the dice (see below). Following this, they were asked to convince themselves that the six-sided dice (which was placed under a black, obscure cup) was not biased in some way by throwing it several times. They were also told explicitly that nobody except themselves could see which numbers they threw, thus, they would have to input the numbers into the computer. To make sure the participants were convinced that nobody could know which numbers they actually threw, it was also explained to them that each participant was instructed to put the dice back under the black cup showing the 1 on the upright position after completing the experiment. Next, the payment rules were explained to them: If they threw a 1, they would get 1 monetary unit (MU) extra payoff for entering a 1 in the computer, for a 2 they would get 2 MUs, and so on (each MU was worth 1 RMB). But if they threw a 6, or rather entered a 6 in the computer, they would get nothing for that round. The participants were asked to throw the dice three times to determine the extra payoff and place the dice with the 1 on the upright position back under the black cup afterwards. While throwing the dice and inputting the numbers in the computer, the rule of payment was always displayed on the screen in form of a table. As nobody other than the participants knew, which numbers they actually threw, they could lie regarding the numbers they threw to maximize their payoff. After the die-in-a-cup paradigm participants were asked to rate how honest they thought other participants would be in the die-in-a-cup paradigm on a 7-point Likert scale (1 = totally dishonest; 7 = totally honest). This was done to assess i) if genetic underpinnings and/or treatment would influence the belief about the honesty of others ii) if differences in the belief about the honesty of others (on group level) would be associated with differences in honesty or lying behaviour (on group level) (see for example [7]).

### Statistical analyses

#### Analysing possible confounding variables

First, the groups of carriers vs. non-carriers of all OXTR haplotypes were compared regarding age and the Honesty-Humility (sub)scales of the HEXACO-PI-R [17] (the HEXACO-PI-R was assessed during the CGBBP and was therefore not influenced by treatment; hence, examining genotype effects across treatment groups is justified). These analyses were implemented using Mann-Whitney U-Tests. Next, differences between PLC and OXT groups were assessed for the same variables as well as the participant’s ratings of honesty of others playing the die-in-a-cup paradigm also using Mann-Whitney U-Tests. Finally, the groups split by PLC vs. OXT treatment and OXTR haplotype carriers vs. non-carriers (2×2 designs) were compared in light of all the previously mentioned possible confounding variables (always comparing 4 groups: PLC, non-carriers vs. PLC, carriers vs. OXT, non-carriers vs. OXT, carriers) using Kruskal-Wallis Tests. Non-parametric analyses were chosen because several of the dependent variables did not fulfil criteria for parametric testing.

#### Analysing the die-in-a-cup paradigm

Analysing the data of the die-in-a-cup paradigm is only possible on group level as individual lies are undetectable with the present experimental set-up. Therefore, the distribution of the reported numbers (numbers inserted in the computer; all three rounds collapsed) was compared with the equal distribution (by chance, each number should have been thrown in 1/6^th^ = 16.67% of the rounds) in the treatment groups (PLC vs. OXT). Additionally, also the effects of treatment (PLC vs. OXT) on the distributions of reported numbers in each round separately were investigated. Lastly, the distributions of the reported numbers for each separate round split by treatment and individual haplotypes (carriers vs. non-carriers; 2×2 designs) were investigated to search for possible interactions. To test statistically for significance of the deviations from the expected equal distribution, Chi^2^-tests were calculated. If Chi^2^-tests revealed significant (p < .05) deviations, the observed frequencies of each individual number were compared with the expected frequency (1/6^th^) using binomial tests (see for example [10] for a similar approach).

## Results

### Possible confounding variables

No significant differences in age or the HEXACO-PI-R [17] Honesty-Humility (sub)scales were observed between the OXTR haplotype groups (for each haplotype comparing carriers vs. non-carriers), which would hold after correction for multiple testing (.05 / 6 = .0083; divided by six because six individual haplotypes on the OXTR gene were investigated) (all p-values > .035).

Mann-Whitney U-Tests revealed also no significant differences between PLC and OXT groups in the possible confounding variables (i. e. age, HEXACO-PI-R Honesty-Humility, and ratings of how honest other participants were thought to be in the die-in-a-cup paradigm – all p-values > .105).

In the groups split by treatment (PLC vs. OXT) as well as carriers and non-carriers of each OXTR haplotype (2×2 designs), Kruskal-Wallis Tests showed no significant difference in the possible confounding variables which would hold after correction for multiple testing (.05 / (6×2) = .0036; divided by 6×2 because six individual haplotypes on the OXTR gene were investigated in two groups each (PLC and OXT) – all p-values > .008). As a result of this, it was decided not to include these variables as confounding variables in further analyses.

### Main effects of treatment on lying behaviour

When investigating our first hypothesis concerning the effects of treatment on lying behaviour, we found that the distribution of the reported numbers (all three rounds collapsed) did not deviate significantly from the equal distribution in the PLC group (Chi^2^(5) = 3.55, p = .616). However, in the OXT group there was a significant deviation from the equal distribution (Chi^2^(5) = 17.27, p = .004). As can be seen in Figure 1, the observed distribution indicates evidence for lying behaviour only in the OXT group. This effect would hold after correction for multiple testing (.05 / 2 = .025; divided by 2 because two groups (PLC vs. OXT) were investigated).

**Figure 1.**
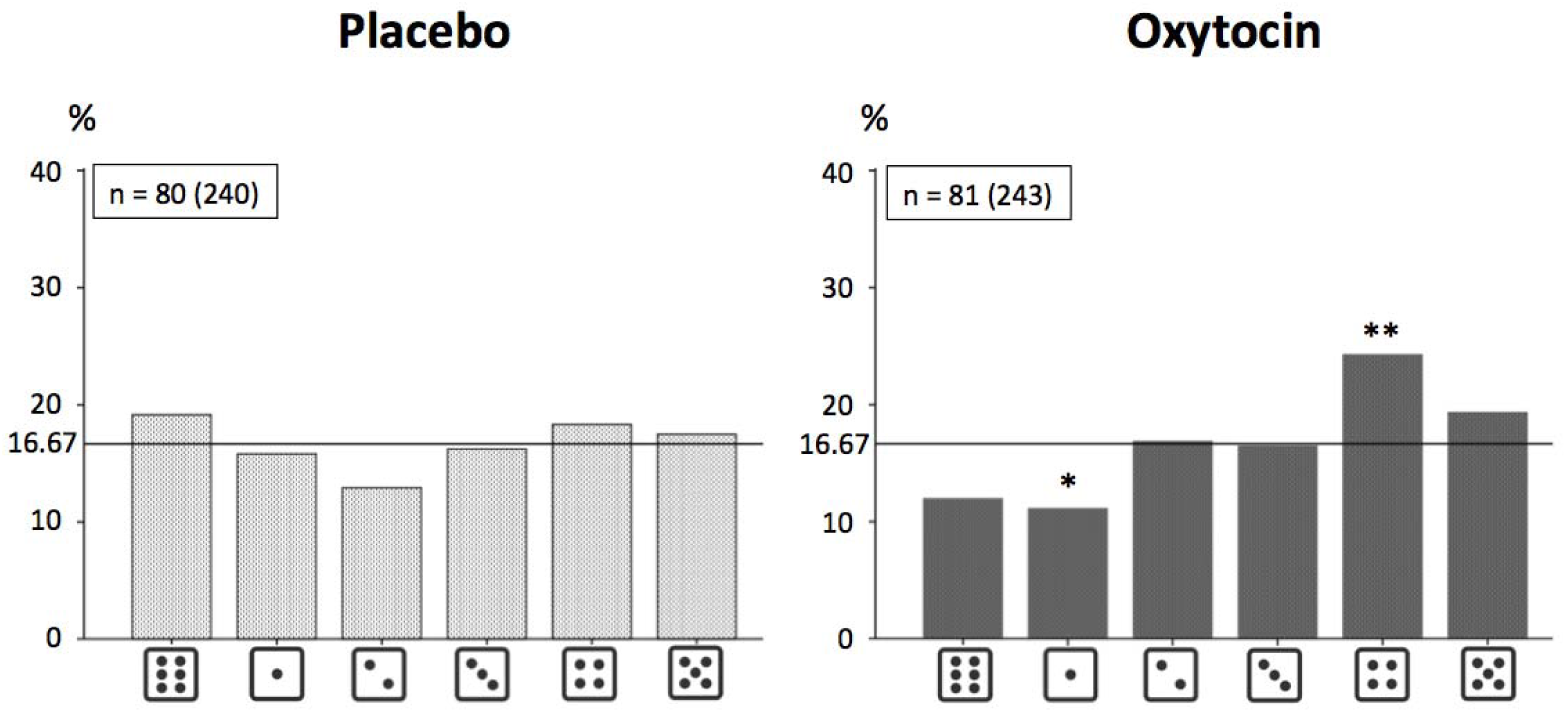
Distributions of numbers reported (in %) in the PLC and OXT groups. Binomial tests were only calculated for the OXT group, where the Chi^2^-test revealed a significant deviation from the equal distribution: *p<.05, **p<.01, ***p<.001 (two-tailed); n =
number of participants in the respective group (number of times the dice was thrown = number of participants in the respective group × 3).

By further investigating each round separately, no significant deviation from the equal distribution in the numbers reported in any round was found in the PLC group (all p-values > .153; for distributions see Supplementary Material). On the other hand, as presented in Figure 2, it was found that lying in the OXT group seemed to be enhanced with each round (in particular with regard to the choice of number 4). The only significant deviation from the equal distribution observed was in the third round (Chi^2^(5) = 22.04, p < .001; all other p-values > .317). This effect would also hold after correction for multiple testing (.05 / (2×3) = .0083; divided by 2×3 because the effects in two groups (PLC vs. OXT) and three rounds were investigated).

**Figure 2.**
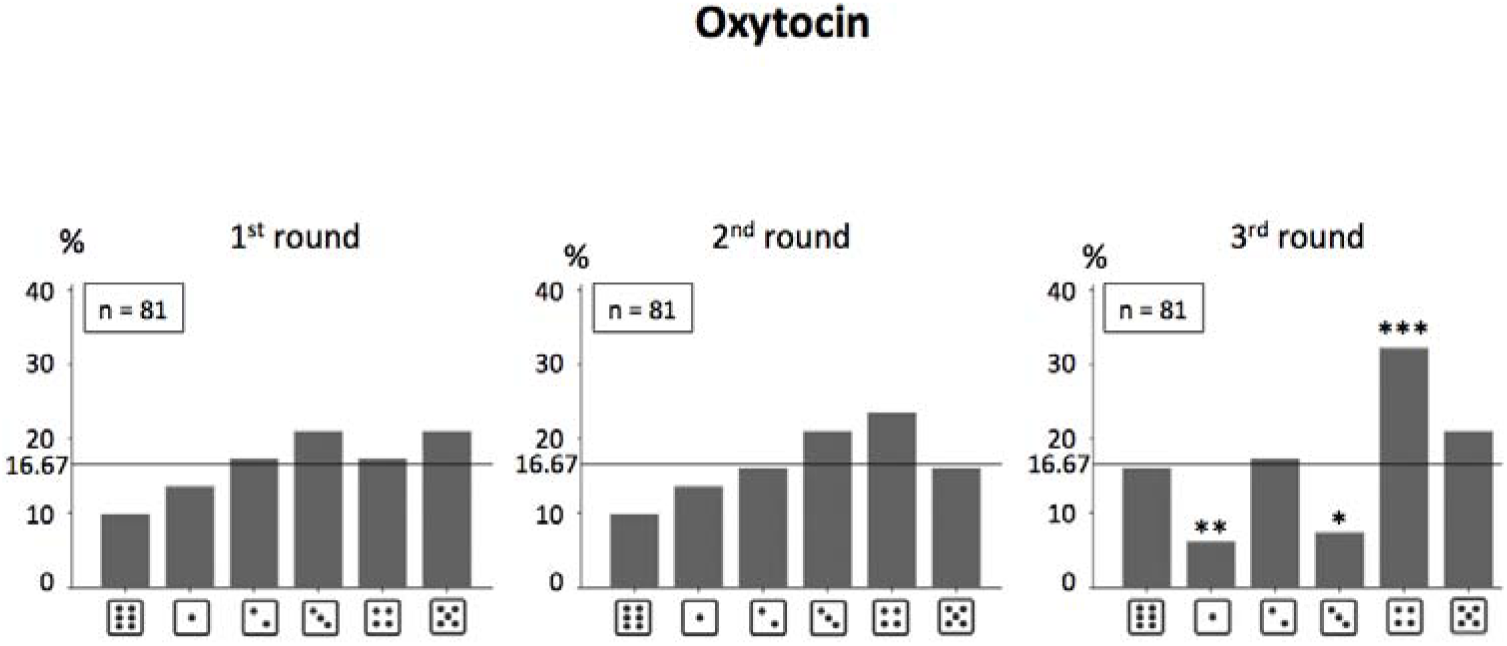
Distributions of numbers reported in the 1^st^, 2^nd^ and 3^rd^ round of the die-in-a-cup paradigm (in %) in the OXT group. Binomial tests were only calculated for the third round as only in this round the Chi^2^-tests revealed a significant deviation from the equal distribution: *p<.05, **p<.01, ***p<.001 (two-tailed), n = number of participants in the respective group.

### Treatment by haplotype interaction effects on lying behaviour

Based on the round-specific effects of OXT on lying behaviour, interaction effects between PLC vs. OXT treatment and individual OXTR haplotypes were analysed on each round separately. In the PLC group, no effect of any individual haplotype (comparing carriers and non-carriers) was found on any round (the respective Chi^2^-Tests revealed p-values > .083). In the OXT group significant results were observed, particularly in the third round (only one significant effect in the second round (p<.05), but none in the first round). However, only two effects in the third round survived a Bonferroni correction procedure (.05 / (6×2×3) = .0014; divided by 6×2×3 because the effects of six individual haplotypes in two groups (PLC vs. OXT) on three rounds were investigated). Results for the third round in the OXT group split by each individual haplotype (carriers vs. non-carriers) are presented in Table 1. In detail, the groups of GCG (rs237887-rs2268491-rs2254298) carriers and GT (rs53576-rs2268498) non-carriers showed a significant deviation from the equal distribution, suggesting that OXT specifically induced lying behaviour in these groups. Therefore, these distributions are also shown in Figures 3 and 4.

**Figure 3.**
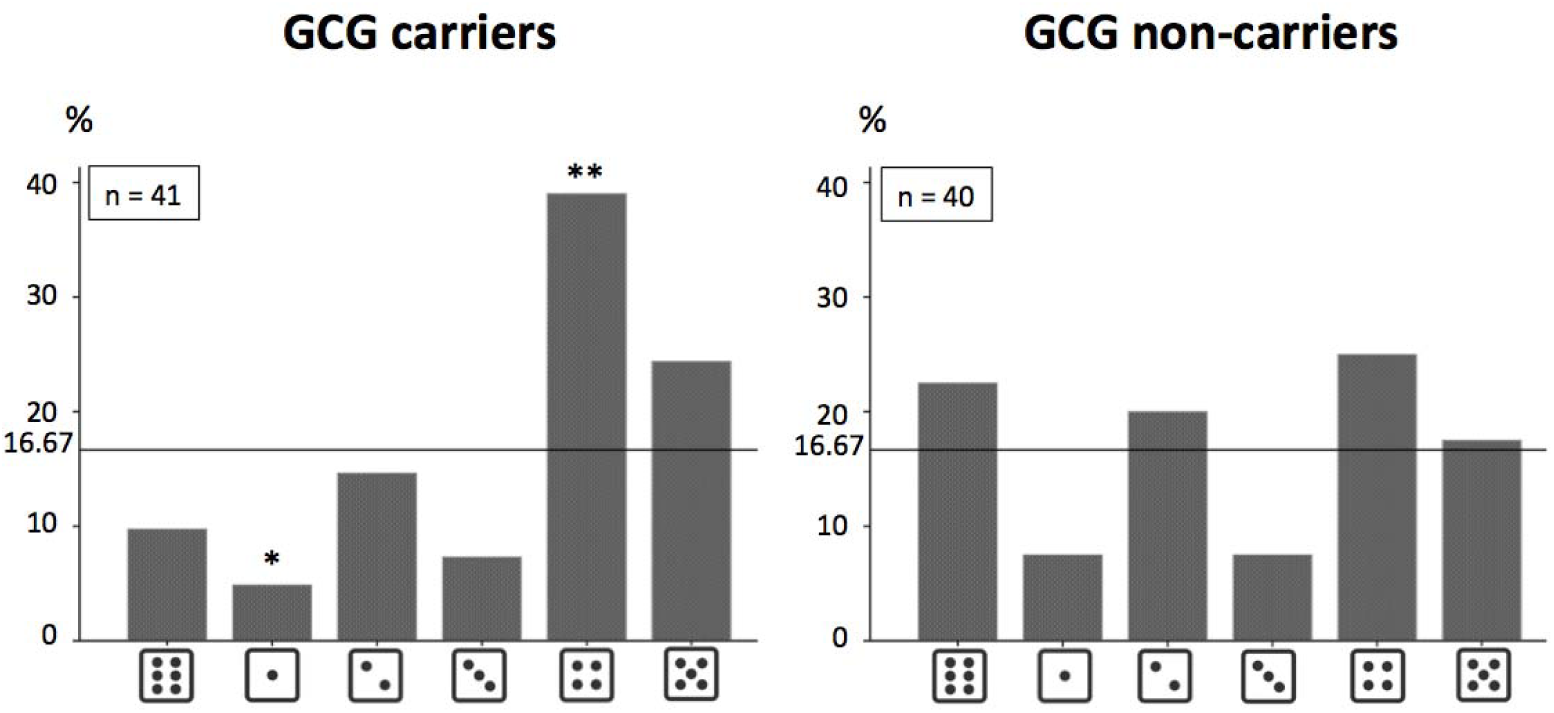
Distributions of numbers reported (in %) in the 3^rd^ round in GCG individual haplotype (rs237887-rs2268491-rs2254298) carriers and non-carriers in the OXT group. Binomial tests were only calculated for the carriers group as only in this group the Chi^2^-test revealed a significant deviation from the equal distribution: *p<.05, **p<.01, ***p<.001 (two-tailed); n = number of participants in the respective group.

**Figure 4.**
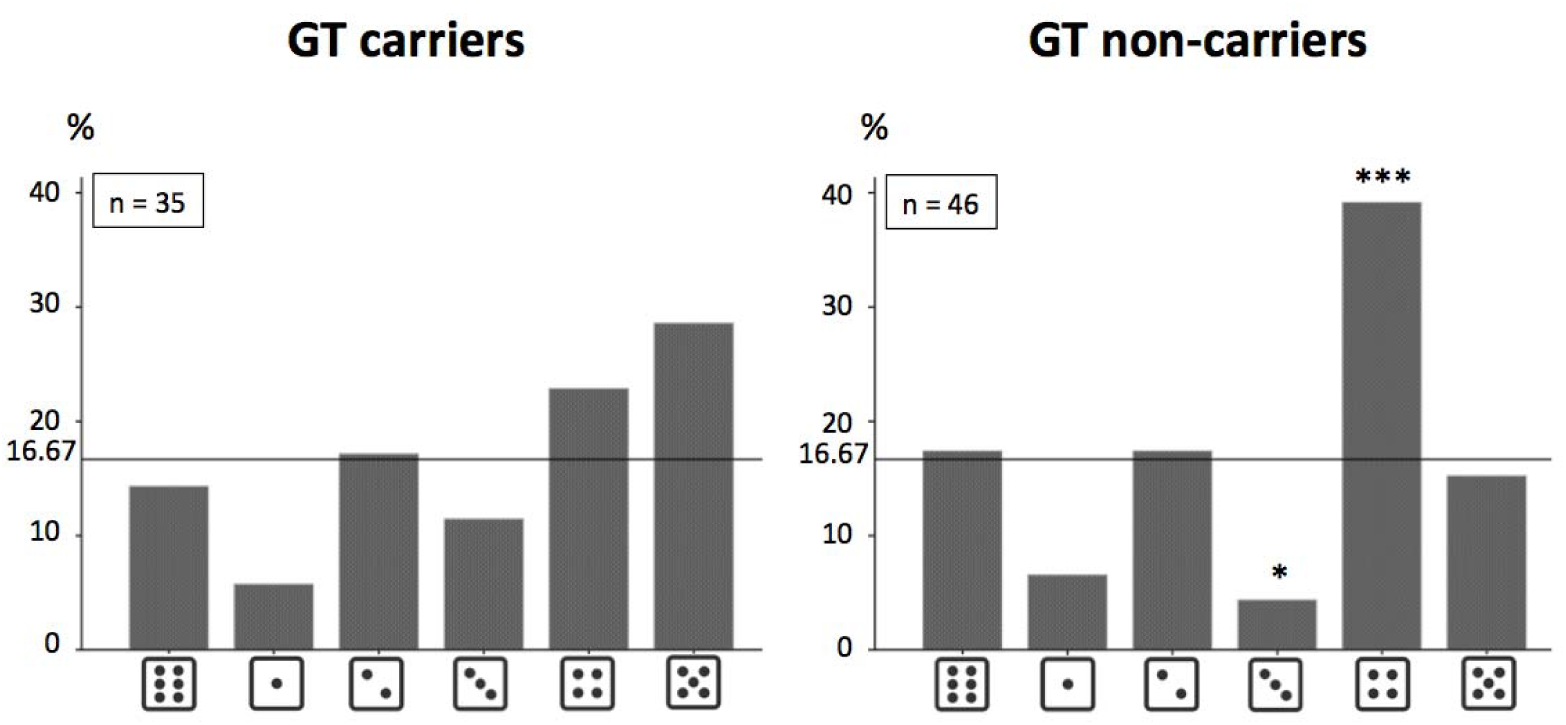
Distributions of numbers reported (in %) in the 3^rd^ round in GT individual haplotype (rs53576-rs2268498) carriers and non-carriers in the OXT group. Binomial tests were only calculated for the non-carriers group as only in this group the Chi^2^-test revealed a significant deviation from the equal distribution: *p<.05, **p<.01, ***p<.001 (two-tailed); n = number of participants in the respective group.

**Table 1.** Statistics for the deviation of the distributions of reported numbers in the third round from the equal distribution in the OXT group for carriers and non-carriers of the OXTR haplotypes of interest

|  | OXT |  |
| --- | --- | --- |
|  | carriers | non-carriers |
| rs237887 - rs2268491 - rs2254298 |  |  |
| GCG | **Chi<sup>2</sup>(5)=20.61,**<br>**p<.001** | Chi<sup>2</sup>(5)=6.80,<br>p=.236 |
| GTA | Chi<sup>2</sup>(5)=8.21,<br>p=.145 | Chi<sup>2</sup>(5)=19.47,<br>p=.002 |
| ACG | Chi<sup>2</sup>(5)=19.56,<br>p=.002 | Chi<sup>2</sup>(5)=3.89,<br>p=.566 |
| rs53576 - rs2268498 |  |  |
| AC | Chi<sup>2</sup>(5)=16.00,<br>p=.007 | Chi<sup>2</sup>(5)=11.62,<br>p=.040 |
| AT | Chi<sup>2</sup>(5)=19.40,<br>p=.002 | Chi<sup>2</sup>(5)=1.62,<br>p=.805 |
| GT | Chi<sup>2</sup>(5)=7.00,<br>p=.221 | **Chi<sup>2</sup>(5)=21.04,**<br>**p<.001** |

## Discussion

The present study sought to investigate potential effects of intranasal OXT administration and OXTR genetics on self-serving lying with a particular focus on possible interactions. In support of our first hypothesis, a significant effect of intranasal OXT treatment on self-serving lying behaviour was observed: for the OXT group lying could be inferred whereas this was not true for the PLC group. Interestingly, in the OXT group lying behaviour was particularly increased in the third and final round of the die-in-a-cup paradigm. This also raises potential methodological issues for future studies in which lying behaviour or other anti-social behaviours are investigated since these may only occur when subjects are given repeated opportunities to display such behaviours. Finally, our results demonstrate important interactions between OXT treatment and OXTR genetics with the intranasal OXT effect on lying behaviour being specific to GCG- (rs237887-rs2268491-rs2254298) carriers and GT- (rs53576-rs2268498) non-carriers. Previous studies have reported that OXT administration only led to increased lying behaviour (compared to PLC) when it benefitted an in-group (including oneself) and in competitive environments, in which participants were concerned that another person would take the money by lying if they did not also do so. But no effects on pure self-serving lying behaviour were observed [6, 7]. This might at first glance seem contradictory to the present results. However, to reconcile the studies and results, it is important to note that the decision about whether to lie can be understood as a consequence of a cost-benefit analysis [3]. On the negative side (against lying) there are the potential costs of getting caught, which were minimal in the present paradigm since detection of lies was not possible. Also, cognitive dissonance and the need to actualize one’s own self-concept after lying (because dishonesty does not match the self-concept of oneself as an honest person) are on the negative side [3]. On the positive side (pro lying), there is the enhancement of the additional payoffs received.

In the OXT administration studies by Shalvi and colleagues and Aydogan et al. [6, 7], coin-toss tasks were used where participants only had two choices: “lie” or “be honest”. In comparison, in the present study participants had more gradations of honesty or lying to choose from. As such, participants could be completely honest, lie to a small extend (e. g. by reporting to have thrown a 2 when actually they threw a 1), or lie to the maximum (by reporting to have thrown a 5 if actually they had thrown a number leading to a much smaller payoff). As can be seen in the distributions of reported numbers (in the groups in which lying could be inferred) in the present study: the 4 rather than the 5 was reported most often. This indicates that participants under OXT did not lie to the maximum, but to a slightly lesser extent. And this can be interpreted as a strategy to maintain a positive self-concept of oneself as an honest person, despite actually lying [3]. Additionally, the findings of the present study are partly in line with results from a cross-cultural study, in which similar patterns (lying in the die-in-a-cup paradigm with regard to the 4 but not 5) in samples from Ho Chi Minh City, Vilnius, Granada and Bogota were found, although in a sample from Shanghai the 5 was reported most often [9]. In the study by Shalvi et al., OXT increased lying behaviour only when subjects could maintain a positive self-concept since their lies benefitted their in-group (as well as themselves) [6]. Similarly, in the study by Aydogan and colleagues [7], OXT also only increased lying behaviour in the context, in which a positive self-concept could be maintained, where lying was in a competitive situation where it avoided an opponent taking the money by lying. Notably, in the present study the effects of OXT on increased lying behaviour were not associated with a belief that other participants would be dishonest when performing the task and a conformity effect therefore cannot explain the present results. In conclusion, in each of the three different studies OXT only seems to increase self-serving lying behaviour in contexts in which participants can lie while still maintaining a positive self-concept There could however also be other possible explanations for the present findings.

Moreover, in the two previous studies where effects of OXT administration on pure self-serving lying behaviour were not detected results are based on an overall response across several rounds. Hence, significant effects of OXT administration on lying behaviour in later rounds might have gone undetected due to non-significant effects in earlier rounds. In the present study, the importance of possible alterations in patterns of lying behaviour as a function of repeated opportunities to lie was confirmed in support of the second hypothesis, namely that there would be round-wise effects of OXT treatment. Only the distribution of the reported numbers in the third round deviated significantly from the equal distribution (although a trend was already visible in the second round) in the OXT group. Revisiting Figure 2, it becomes obvious on a descriptive level that the number of 4 inserted into the computer constantly rises over the three rounds. A possible explanation is that with increasing number of rounds without (negative) consequences of the behaviour, certainty that lying would not be detected and/or punished might be increased and participants under OXT might therefore tend to lie more. However, with the present study design we cannot test this directly and other explanations are also possible. We did not, for example, obtain self-reports on how certain a person was that lying would go undetected across the three rounds, because doing so might have altered behaviour during the task.

At last, interactions of OXT administration and OXTR genetics were found. Other than rs2268498, the OXTR SNPs under investigation are placed in an intronic region of the gene and their functionality on a biochemical level is unknown. Thus, it is not possible to directly conclude potential functional differences (i. e. in oxytocin binding, or receptor density, or structure or functioning of the receptors in the brain / of the brain) between the groups of certain individual haplotype carriers vs. non-carriers. However, since no overall differences were observed between the samples (split by treatment and/or individual haplotypes) in stable personality variables (HEXACO-PI-R Honesty-Humility (sub)scales), and as lying did not occur in any (sub-)group under the influence of PLC but only for specific individual haplotype groups under the influence of OXT, it is reasonable to conclude that observed effects are indeed caused by an interaction of OXT treatment and OXTR genetics. This supports the assumption that OXT effects are highly specific and OXT sensitivity depends on genetic predispositions and additionally underlines the importance of assessing genetic moderators in OXT administration studies [14-16; 19, 20].

Several limitations of the present study should also be acknowledged. Firstly, only males were investigated and since sex-dependent effects of OXT have been reported by previous studies [21, 22] it is possible that OXT effects on self-serving lying might also be different in females. Secondly, rather small sample sizes in some of the subgroups could have influenced our ability to detect effects. However, by grouping subjects into carriers vs. non-carriers of individual haplotypes this problem was minimized and it is unlikely that the present results are an artefact of low sample size (please see Figures 3 and 4 which show that large and nearly equal sized groups were tested). Finally, it is likely that there are additional genes and polymorphisms, which could influence the effects of intranasally applied OXT (e. g. polymorphisms in the CD38 (Cluster of Differentiation 38) gene or in the AVPR1a (arginine-vasopressin receptor 1a) gene (see for example [23-26]). For the present study, however, it was decided to only investigate the most well established candidate gene polymorphisms of the oxytocinergic system with respect to lying, which have previously been associated with social cognition and relevant behaviours and/or have a known biochemical function (e. g. [27-30]).

In conclusion, the findings of the present study demonstrate that OXT plays an important role in self-serving lying and thereon potentially also other (anti-)social behaviours. Moreover, these effects of intranasal OXT administration are highly specific and situational factors, such as the repeated chance to lie undetected, and genetic underpinnings of the OXTR gene moderate them. Therefore, to gain a deeper understanding of the effects of OXT treatment, and to help interpret heterogeneous results in the literature it may help if future OXT administration studies also take into account the genotypes of participants.

## Funding and Disclosure

### Author Contributions

C.S. and C.M. planned the design of the present study. B.B. and K.M.K. gave helpful advice to improve the study design. C.S. and R.L. programmed the paradigms and assessed the data (genetic as well as experimental). C.S. and R.L. conducted the genetic analyses. C.S. wrote the manuscript and carried out the statistical analyses. C.M., B.B. and K.M.K. worked over the manuscript. All authors approved the final version of the manuscript and agreed submission.

## Acknowledgements

None.

